# BWA-mem is not the best aligner for ancient DNA short reads

**DOI:** 10.1101/2021.08.02.454401

**Authors:** Adrien Oliva, Raymond Tobler, Bastien Llamas, Yassine Souilmi

**Author notes:** Corresponding Author: Y.S.

## Abstract

Xu and colleagues (Xu et al., 2021) recently suggested a new parameterisation of *BWA-mem* (Li, 2013) as an alternative to the current standard *BWA-aln* (Li and Durbin, 2009) to process ancient DNA sequencing data. The authors tested several combinations of the *-k* and -r parameters to optimise *BWA-mem*’s performance with degraded and contaminated ancient DNA samples. They report that using *BWA-mem* with −*k* 19 −*r* 2.5 parameters results in a mapping efficiency comparable to *BWA-aln* with −*I* 1024 −*n* 0.03 (i.e. a derivation of the standard parameters used in ancient DNA studies; (Schubert et al., 2012)), while achieving significantly faster run times.

We recently performed a systematic benchmark of four mapping software (i.e. *BWA-aln*, *BWA-mem*, *NovoAlign* (http://www.novocraft.com/products/novoalign), and *Bowtie2* (Langmead and Salzberg, 2012) for ancient DNA sequencing data and quantified their precision, accuracy, specificity, and impact on reference bias (Oliva et al., 2021). Notably, while multiple parameterisations were tested for *BWA-aln*, *NovoAlign*, and *Bowtie2*, we only tested *BWA-mem* with default parameters.

Here, we use the alignment performance metrics from Oliva et al. to directly compare the recommended *BWA-mem* parameterisation reported in Xu et al. with the best performing alignment methods determined in the Oliva et al. benchmarks, and we make recommendations based on the results.

## Methods

We investigated the alignment performance of the parameterisation recommended by Xu et al., i.e. −*k* 19 and −*r* 2.5 (hereafter called BWA9) against several of the best performing strategies identified in Oliva et al. (namely, BWA1, BWA2, BWA8, Novo1IUPAC, Novo2IUPAC, and Novo2, see Table 1 for parameter settings).

**Table 1.**
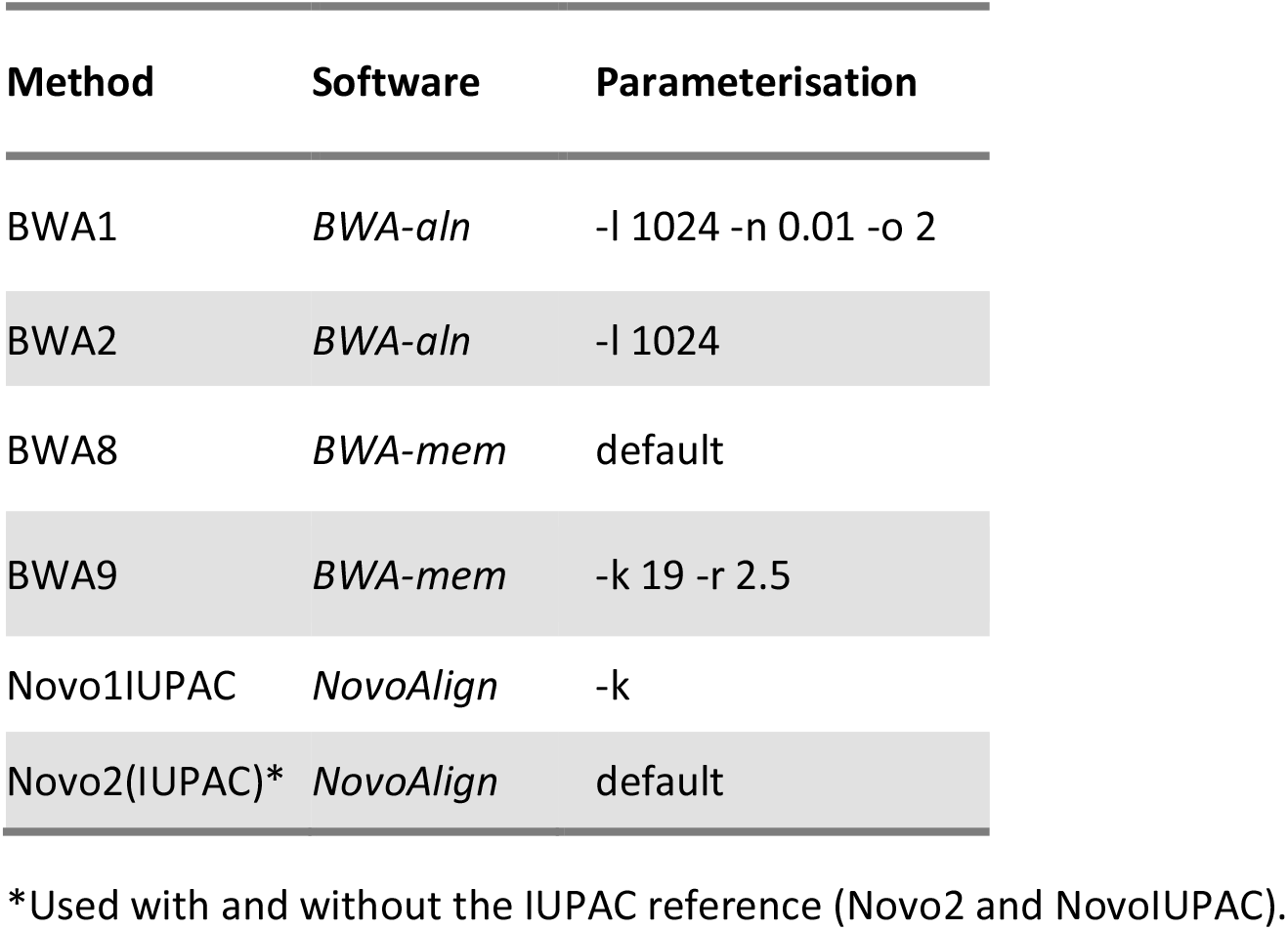
Different alignment parameterisations tested.

| Method | Software | Parameterisation |
| --- | --- | --- |
| BWA1 | *BWA-aln* | -l 1024 -n 0.01 -o 2 |
| BWA2 | *BWA-aln* | -l 1024 |
| BWA8 | *BWA-mem* | default |
| BWA9 | *BWA-mem* | -k 19 -r 2.5 |
| Novo1IUPAC | *NovoAlign* | -k |
| Novo2(IUPAC)* | *NovoAlign* | default |

\*Used with and without the IUPAC reference (Novo2 and NovoIUPAC).

Following the analytical framework of Oliva et al., our benchmark is based on simulated reads (including fragmentation, damage, and sequencing errors typical for ancient DNA samples; see (Oliva et al., 2021)) that were generated for each of the following three samples from the 1000 Genome Project (1000 Genomes Project Consortium et al., 2015) dataset, each coming from a distinct population, and were aligned to reference genome GRCh37:

- Individual *HG00119* from the British in England and Scotland population; GBR; labelled Europe in this study.
- Individual *NA19471* from the Luhya population in Webuye, Kenya; LWK; labelled Africa in this study.
- *Individual HG00513* from the Han Chinese population in Beijing, China; CHB; labelled East Asia in this study.

In addition to quantifying read alignment precision (i.e. the proportion of correctly aligned reads relative to all aligned reads) and proportion of aligned reads (i.e. the fraction of aligned reads relative to the total number of simulated reads) for each strategy, we tested the specificity (i.e., the fraction of unmapped reads) of these strategies for two sets of potential contaminants—i.e. bacterial and dog reads—that were also used in Oliva et al., 2021.

## Results

BWA9 had a slight improvement in the proportion of total reads aligned relative to *BWA-mem* using default settings (BWA8), but this came at the cost of consistently lower precision (Figure 1). These precision differences are particularly accentuated for reads between 30 to 60bp, the range of read lengths that is typical of ancient DNA. As demonstrated here and in more detail in our recent alignment software benchmark (Oliva et al., 2021), *BWA-aln* (BWA1 and BWA2) is the most precise alignment method amongst the tested strategies, having moderately higher precision relative to *BWA-mem* for shorter reads while mapping a much higher percentage of reads overall (Oliva et al., 2021; van der Valk et al., 2021).

**Figure 1.**
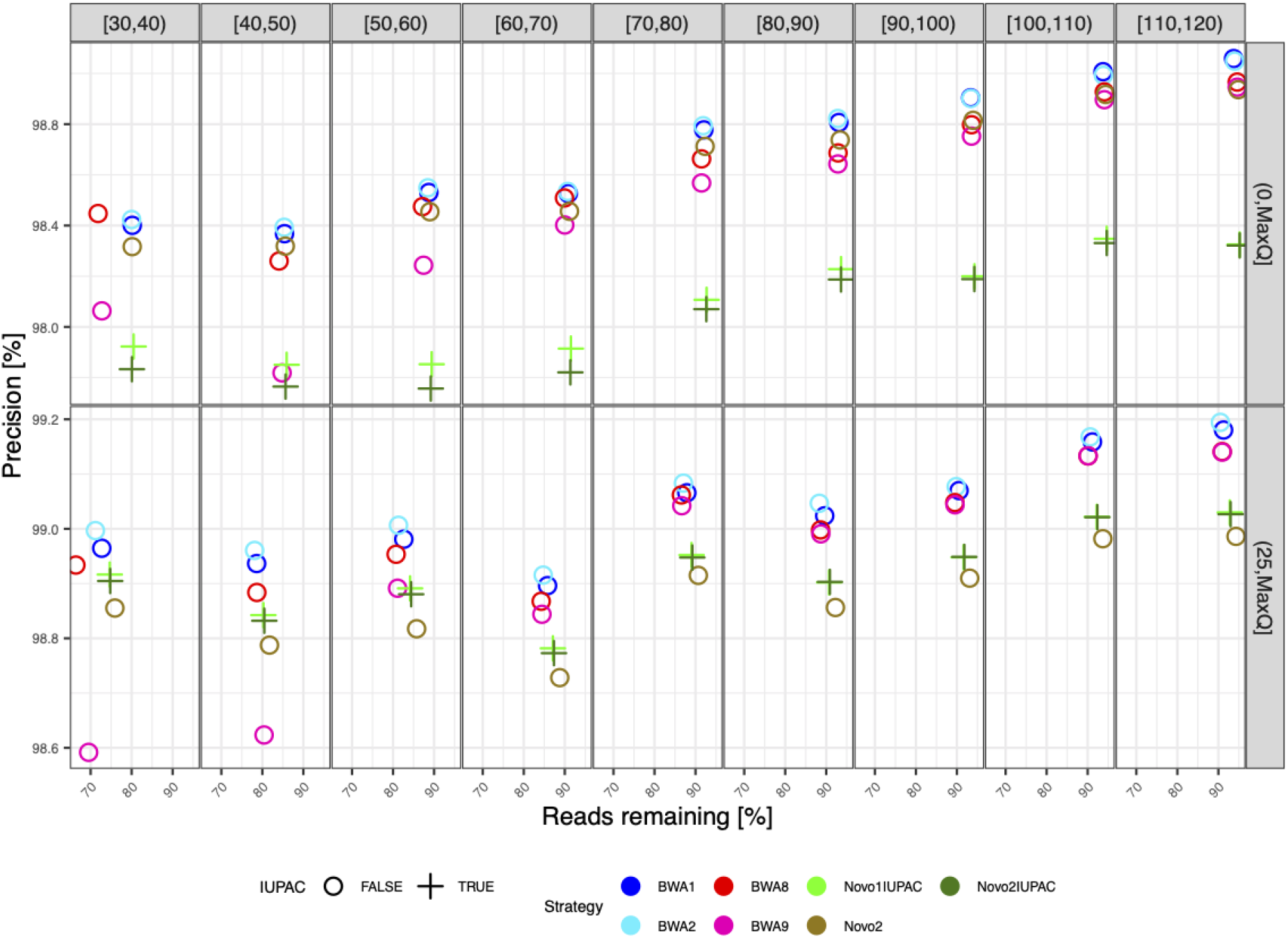
Alignment precision relative to read length and mapping quality for the simulated East Asian sample. Results are shown for seven parameterisations of four different alignment software, including an IUPAC reference-based alignment for a subset of the *NovoAlign* parameterisations (see key). BWA9 is the *BWA-mem* strategy recommended by Xu et al., 2021, with parameter details for the other strategies provided in Table 1. The panels in each row show results after applying the specific mapping quality filter, which results in the removal of all reads below the required mapping quality. Results were similar for the simulated European and African samples and are shown in Appendix Figures 1 and 2, respectively.

When comparing specificity against potential contaminants, BWA9 has a near identical specificity to the default *BWA-mem* parameterisation (BWA8) for dog reads, and slightly poorer specificity when testing with bacterial reads, but both parameterisations perform considerably worse than the tested *NovoAlign* (Novo1IUPAC, Novo2IUPAC, and Novo2) and *BWA-aln* (BWA1 and BWA2) strategies for dog reads (Figure 2).

**Figure 2.**
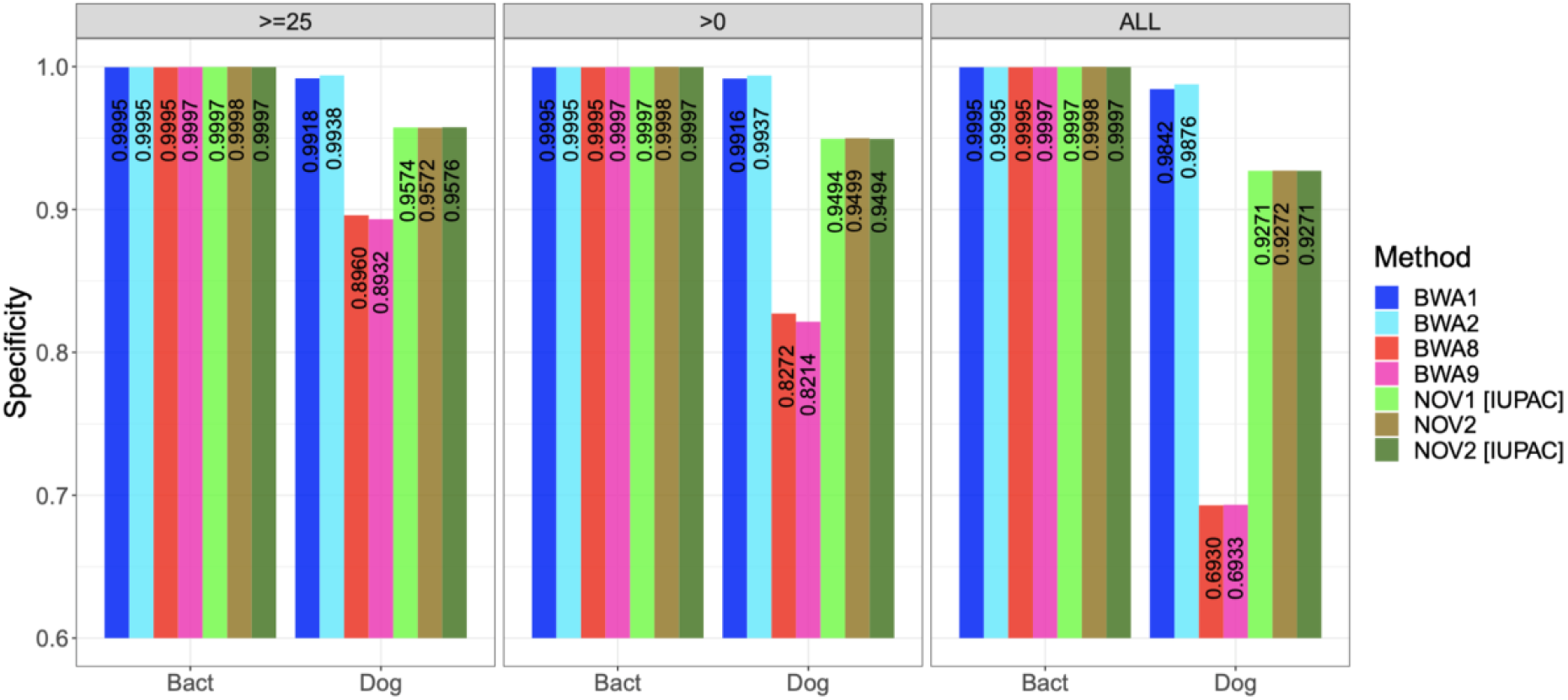
Specificity of all tested alignment methods. Bacterial and dog reads were aligned to the GRCh37 reference using the seven tested parameterisations of four different alignment software, including an IUPAC reference-based alignment for a subset of the *NovoAlign* parameterisations (see key). The specificity corresponds to the number of unmapped reads, with higher values being better.

Finally, comparing running times of the two *BWA-mem* parameterisations for each of the three simulated human datasets showed that BWA9 is slightly quicker than BWA8 (Figure 3), confirming the results of ref. (Xu et al., 2021).

**Figure 3:**
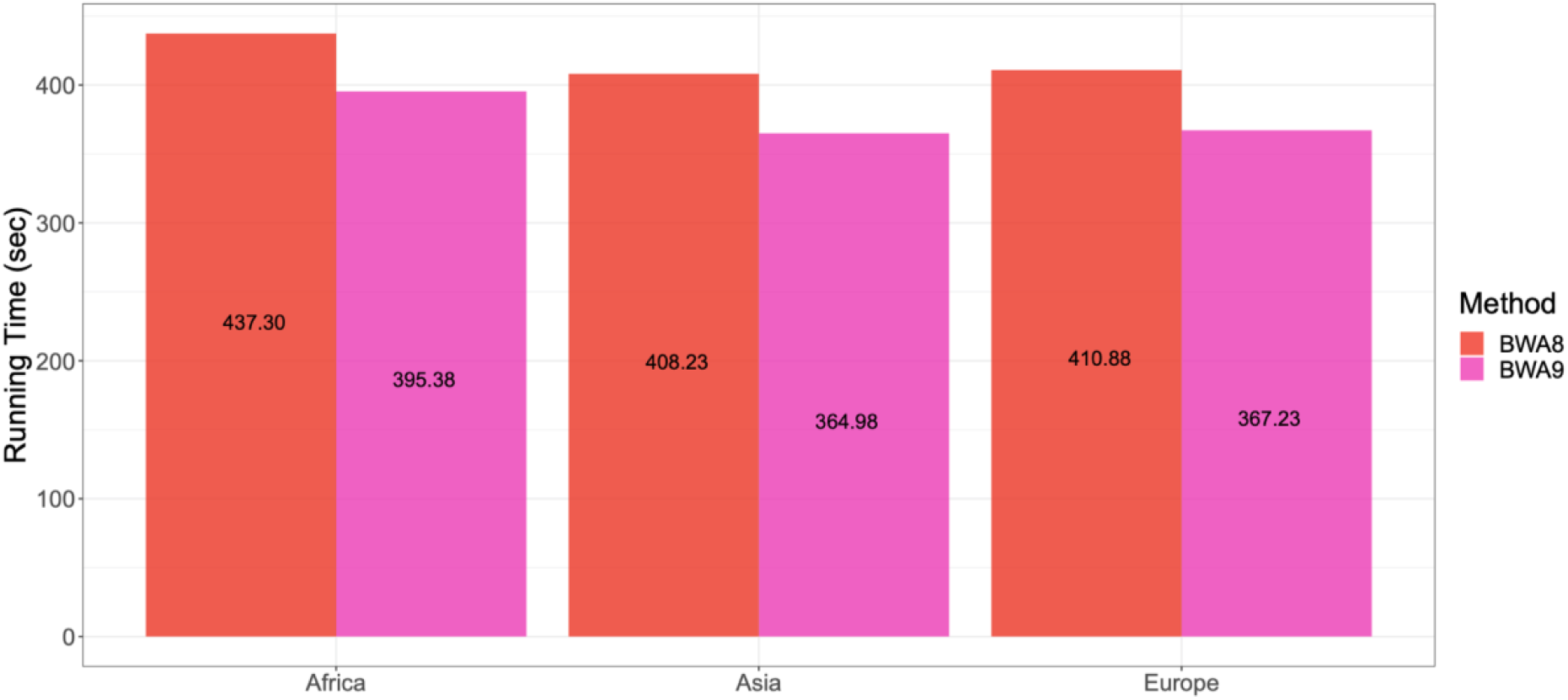
Execution time for each of the *BWA-mem* strategies. The execution time (walltime) in seconds of BWA8 (default parameters) and BWA9 (Xu et al. parameterisation; −k 19 −r 2.5) based on 1.5 million simulated reads.

## Conclusion

Xu et al. report that *BWA-mem* produces alignment results that are comparable to a derivation of the widely used BWA-aln in the ancient DNA field. Consequently, they recommend the use of a specific non-default *BWA-mem* parameterisation for ancient DNA studies because of its superior runtime and specificity relative to *BWA-aln*. However, we find that this parameterisation actually decreases alignment precision relative to *BWA-mem* using default settings for sequencing reads shorter than 70 bases, which are particularly abundant in ancient DNA samples. Moreover, *BWA-mem* is consistently outperformed by *BWA-aln* under the tested parameterisations for both precision and the proportion of reads mapped, and also had greatly improved specificity when the DNA contamination came from a phylogenetically related organism (i.e. a dog in the present study). Crucially, Oliva et al. have demonstrated that improvements in these alignment metrics are also complemented by a reduction in reference genome bias—an alignment-related bias that can inflate false positives and is particularly problematic for ancient DNA studies.

Accordingly, despite having improved run times, we do not recommend that *BWA-mem* is prioritised over *BWA-aln* for research using short reads—such as ancient DNA, cell-free DNA, and forensic research fields. If run time is an issue for researchers, we recommend the use of *NovoAlign* using the free default parameterisation, so long as an appropriate IUPAC reference can be generated. Readers interested in more detailed discussion of these issues are directed to refs. (Oliva et al., 2021; Poullet and Orlando, 2020; Schubert et al., 2012; van der Valk et al., 2021) for recent benchmarks of different alignment strategies using short reads.

## Acknowledgments

We thank the Novocraft Technologies team for providing access to their proprietary *NovoAlign*.

## Data Accessibility

The scripts used to create the used datasets in this study are available in the github repository at: https://github.com/AdrienOliva/Benchmark-aDNA-Mapping.

## Appendix

**Appendix Figure 1:**
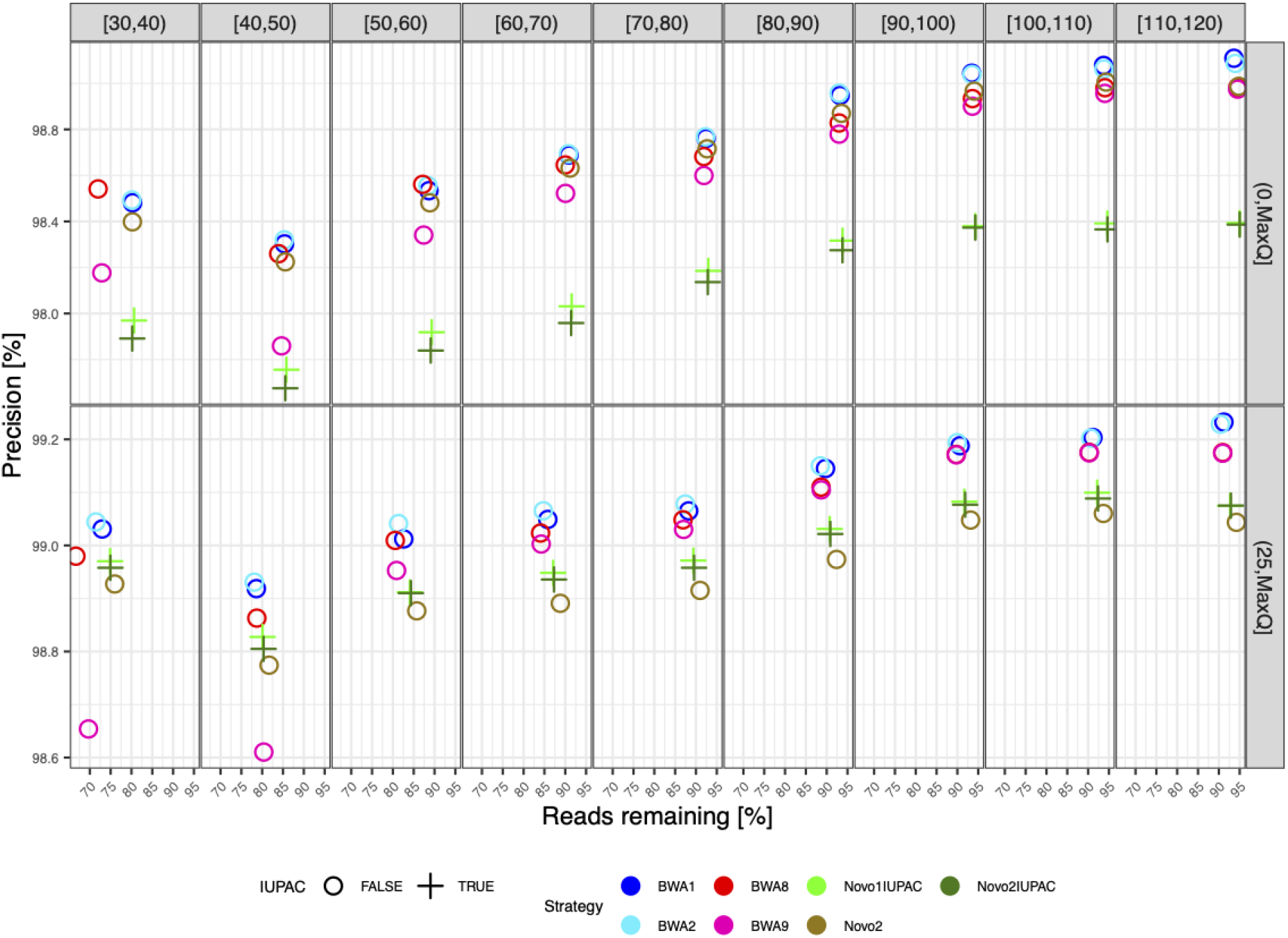
Alignment precision relative to read length and mapping quality for the simulated European sample. See Figure 1.

**Appendix Figure 2:**
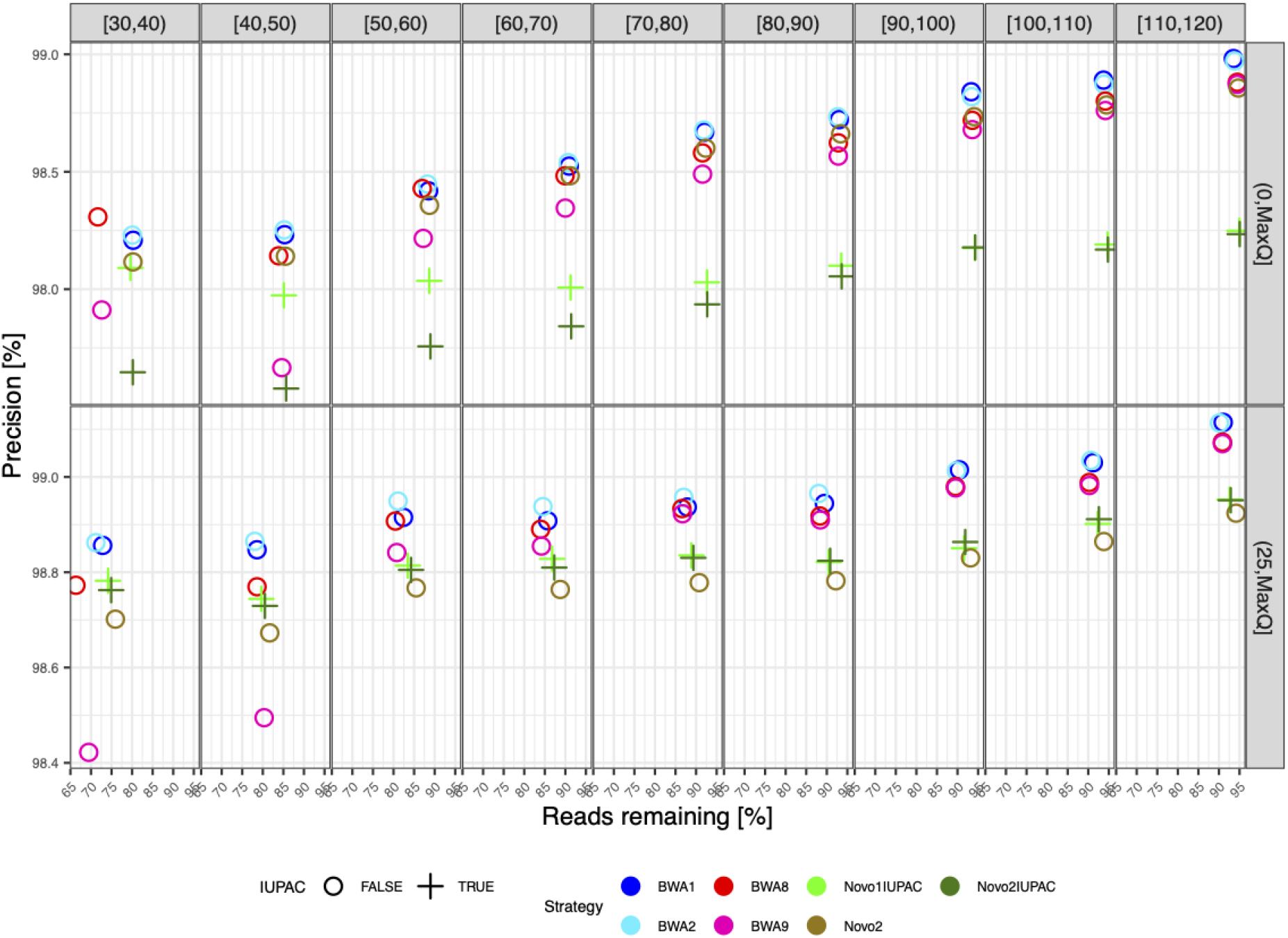
Alignment precision relative to read length and mapping quality for the simulated African sample. See Figure 1.s

